# Dynamics of multiple interacting excitatory and inhibitory populations with delays

**DOI:** 10.1101/360479

**Authors:** Christopher M. Kim, Ulrich Egert, Arvind Kumar

## Abstract

A network consisting of excitatory and inhibitory (EI) neurons is a canonical model for understanding cortical network activity. In this study, we extend the EI network model and investigate how its dynamical landscape can be enriched when it interacts with another excitatory (E) population with transmission delays. Through analysis and simulations of a rate model and a spiking network model, we study the transition from stationary to oscillatory states by analyzing the Hopf bifurcation structure in terms of two network parameters: 1) transmission delay between the EI subnetwork and the E population and 2) inhibitory couplings that induce oscillatory activity in the EI subnetwork. We find that the critical coupling strength can strongly modulate as a function of transmission delay, and consequently the stationary state is interwoven intricately with oscillatory states generating different frequency modes. This leads to the emergence of an isolated stationary state surrounded by multiple oscillatory states and cross-frequency coupling develops at the bifurcation points. We identify the possible network motifs that induce oscillations and examine how multiple oscillatory states come together to enrich the dynamical landscape.

## I. INTRODUCTION

The brain is organized as a network of highly specialized subnetworks. Each of the subnetworks consists of a large number of excitatory and inhibitory neurons communicating via spikes. Randomly connected networks of excitatory and inhibitory neurons have been a popular and useful model to study the dynamical states and information processing in local networks of the brain. Previous work has demonstrated that balance of excitation and inhibition (EI-balance) is a crucial variable that determines two qualitatively different states of global network activity. When excitation and inhibition are balanced, cancellation of excitatory and inhibitory synaptic inputs to a neuron leads to asynchronous and nearly Poisson type spiking [1, 2]. A mismatch between excitation and inhibition (in amplitude or timing) results in oscillatory states [3–5], in which the population firing rate oscillates while individual neurons spike irregularly. Both network states are considered to play important roles in cortical processing; the asynchronous activity of the balanced state provides a suitable substrate to perform complex computations [6, 7], balanced amplification [8, 9] and propagation of rate and time coded signals [10], and oscillatory rhythms play a crucial role in selective routing information across multiple brain areas [11–14].

Besides the EI-balance, spike propagation time delays introduce various complex effects on the network activity dynamics. For instance, delays may destabilize the balanced state of spiking networks [3–5], enrich the bifurcation structure of spatially-extended neural field models by introducing novel dynamical states [15, 16] and gate the propagation of spiking activity [17]. Moreover, by suitably tuning the delays oscillations can be enhanced and suppressed [18–20].

Here we go beyond the standard two population (one excitatory and one inhibitory) model of local cortical networks and investigate how the addition of one more excitatory population alters the dynamical landscape of the standard excitatory-inhibitory (EI) spiking networks composed of balanced and oscillatory states. The third excitatory population is coupled to standard EI network with a longer delay as compared to the delays within in the EI network. Thus, this model allows us to understand how long and short delays interact to shape the critical coupling strength between excitatory-inhibitory or inhibitory-inhibitory neurons that generate network-wide oscillations [3–5].

Through analysis and simulations of the three-population network models, we show that the balanced state can be interwoven intricately with multiple oscillatory states when the EI network is strongly coupled to a third excitatory population with delays. Such dynamical landscapes naturally give rise to cross-frequency oscillations in parameter regime where multiple oscillatory states merge. Our study demonstrates the rich dynamic repertoire of interacting sub-networks in the presence of delays and paves the way to study a network of many sub-networks.

## II. NETWORK MODEL

The network consisted of two excitatory (*E*_1_, *E*_2_) and one inhibitory (*I*_3_) populations. The *E*_2_*I*_3_ subnetwork was the standard EI network [3] which was reciprocally connected to the *E*_1_ population (Fig. 1). We did not include recurrent excitatory connections within *E*_1_ and *E*_2_ because we focused on oscillatory network activity generated by the excitatory-inhibitory and inhibitory-inhibitory couplings. We refer to the connectivity parameters between *E*_1_ and *E*_2_*I*_3_ “lateral”, and those within the *E*_2_*I*_3_ subnetwork “local”. The population *E*_1_ was considered to be located at a farther distance; therefore, the transmission delay *D* between *E*_1_ and *E*_2_*I*_3_ was larger than the transmission delay *d* within the *E*_2_*I*_3_ subnetwork. In this work we investigated the effect of long transmission delays and inhibitory connections from *I*_3_ to *E*_1_, *E*_2_ and itself on network oscillations.

**FIG. 1.**
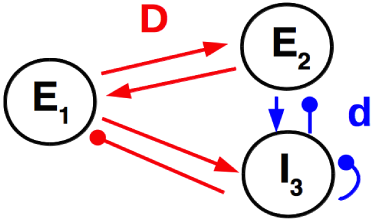
The connectivity structure of *E*_1_*E*_2_*I*_3_ network.

To study the network dynamics analytically, we considered a rate model that describes the firing rate dynamics of three populations and compared the results with numerical simulations of a comparable network with spiking neurons.

### Firing rate model

The rate model was described as a set of delay differential equations,

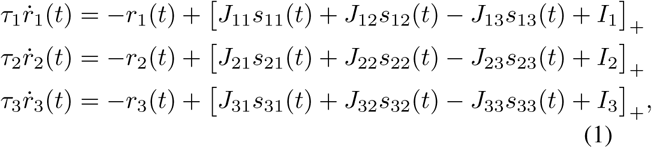

where *J_ab_* is the coupling strength of connection from population *b* to population *a, I_a_* is an external input, and the activation function [*x*]_+_ = *x* if *x* > 0 and = 0 otherwise. We let *J*_11_ = *J*_22_ = 0 because there were no recurrent excitatory connections. The dynamics of synaptic current from *b* to *a* obeyed

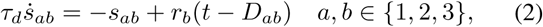

where *τ_d_* is a decay time constant and *D_ab_* is a transmission delay from *b* to *a*.

The connections between *E*_1_ and *E*_2_*I*_3_ had a transmission delay, *D*, and the connections within the *E*_2_*I*_3_ subnetwork had a transmission delay, *d*; *D* = *D*_12_ = *D*_21_ = *D*_13_ = *D*_31_ and *d* = *D*_23_ = *D*_32_ = *D*_33_.

### Network model with spiking neurons

For the spiking network model, we considered a network of randomly connected leaky integrate-and-fire (LIF) neurons where *E*_1_, *E*_2_ and *I*_3_ population consists of *N*_1_ = *N*/2, *N*_2_ = *N* and *N*_3_ = *N*/4 neurons (*N* = 10, 000), respectively. The membrane potential of neuron *i* in population *a* obeyed

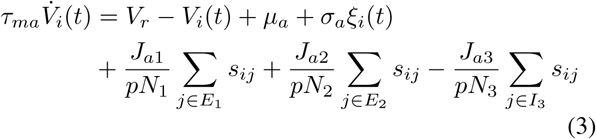

and elicited an action potential when *V_i_* reached a threshold *V_th_*. Here, *τ_ma_* is a membrane time constant, *V_r_* is a resting potential, *J_ab_* is total postsynaptic potential of synaptic connections from population *b* to population *a, µ_a_* is an external input, and *σ_a_ξ_i_* is Gaussian white noise with mean zero and variance 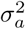.

Every neuron received the same number *pN_a_* of recurrent synaptic inputs from randomly selected neurons in populations *a* = 1, 2, 3. The strength of individual synapses from neuron *i* in population *a* to neuron *j* in population *b* was given by *J_ab_*/(*pN_b_*)

The synaptic current *s_ij_* decayed exponentially upon receiving a spike from a presynaptic neuron *j* in population *b*.

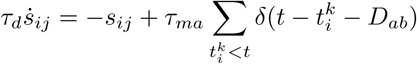

where 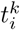 is the spike-time of presynaptic neurons, and *D_ab_* is the transmission delay from neurons in population *b* to neurons in population *a*. As in the firing rate model, the connections between *E*_1_ and *E*_2_*I*_3_ had a transmission delay, *D*, and the connections within the *E*_2_*I*_3_ subnetwork had a transmission delay, *d*.

Following the previous studies [3, 5, 6], we estimated the steady-state firing rate using the Fokker-Planck approach, which meant solving a system of three nonlinear equations in a self-consistent manner.

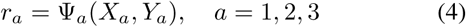

where

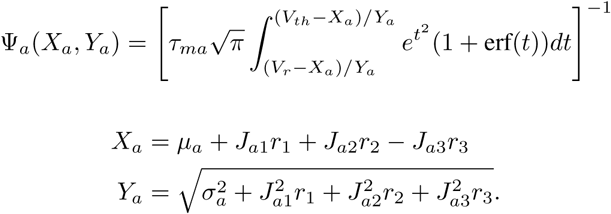

Here, *r_a_* is the population-averaged firing rate, *X_a_* is the mean synaptic input, and 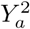 is the variance of total synaptic input to a neuron.

In the following network simulations, we adjusted the mean (*µ_a_*) and variance 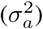 of external inputs to obtain the same steady-state firing rate (*r_e_* = 5Hz, *r_i_* = 10Hz) for different network configurations. The external inputs (*I_a_*) to the rate model was adjusted similarly to maintain same *r_a_* across simulations. Firing rate equations (Eq. 1) were solved using Matlab’s delay differential equation solver, dde23. The simulation of networks with spiking neurons was performed using the NEST simulation tool [21].

## III. DYNAMICS OF OSCILLATIONS IN THE THREE POPULATION MODEL

### A. Linear stability of the steady-state

To characterize how the lateral delay *D* affects the emergence of oscillatory activity within the *E*_2_*I*_3_ subnetwork, we performed linear stability analysis of the steady-state of the rate model and the spiking network model. We added a small perturbation term to the steady-state firing rate such that *r_a_*(*t*) = *r_a_*_0_ + *δr_a_* · *e^λ t^*. The rate perturbation induced a perturbation in the synaptic current:

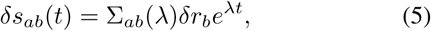

where the synaptic kernel (Eq. 2) in frequency domain is given by

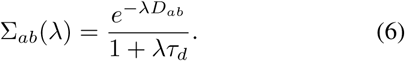

From Eq. 5, we obtained the perturbation of total input to population *a*

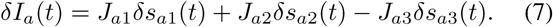

Finally, the new output rate of the network in response to the input rate perturbation is given by

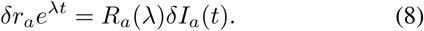

Similar to the synapse, neuron population also acts like a frequency filter which can written as, for the rate model,

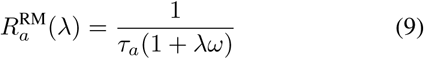

and, for LIF neurons, 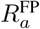 can be either calculated numerically using the threshold integration method [22] or by hypergeometric functions [3, 5].

Combining Eqs. 5, 7 and 8, we obtained a system of three linear equations in *δr_a_, a* = 1, 2, 3, which has non-trivial solutions when the determinant of following matrix is zero:

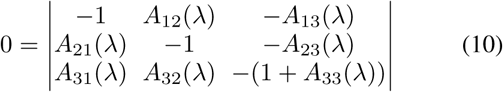

where *A_ab_*(*λ*) = *J_ab_* Σ_*ab*_(*λ*)*R_a_*(*λ*). Rearranging Eq. (10), we obtained

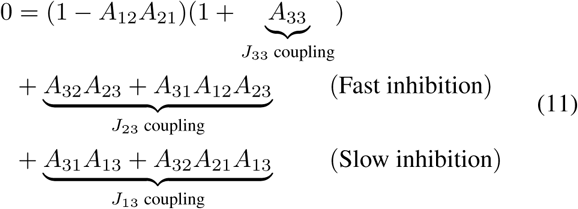

Eq. 11 describes network motifs in the *E*_1_*E*_2_*I*_3_ model that can induce network oscillations. The two first lines of Eq. 11 consist of motifs that receive fast inhibition with short delay *d* via two different pathways. The local inhibitory coupling *J*_33_ is responsible for generating oscillatory activity via the monosynaptic *A*_3_*A*_3_ loop, whereas the inhibitory coupling *J*_23_ is responsible for generating oscillatory activity via disynaptic *A*_32_*A*_23_ and trisynaptic *A*_31_*A*_12_*A*_23_ loops. The third line of Eq. 11 consists of network motifs that receive slow inhibition. In this case, the inhibitory coupling *J*_13_ is responsible for generating oscillatory activity via disynaptic *A*_31_*A*_13_ and trisynaptic *A*_32_*A*_21_*A*_13_ loops with long delay *D*. In the following we systematically investigate how the slow and fast inhibitory pathways interact to shape the oscillatory dynamics of the three population network.

### B. Transition to oscillatory states

To obtain analytical estimates of a Hopf bifurcation, which marks the transition from a steady state to an oscillatory state, we substituted *λ* = *iω* into Eq. 11 to find bifurcation points as a function of *D* and *J*_33_ (or *D* and *J*_23_). A similar calculation was performed in [5]. We denoted the amplitude and the phase of the population response function *R_a_*(*iω*) as *H_a_* and *ϕ_a_*, respectively, i.e. *R_a_*(*iω*) = *H_a_*(*ω*) exp(−*iϕ_a_*(*ω*)). The amplitude and the phase of synaptic kernel ∑_*ab*_(*iω*) were denoted as *H_s_* and *ϕ_s_*, respectively, i.e. ∑_*ab*_ = *H_s_*(*ω*) exp(−*iϕ_s_*(*ω*) −*iD_ab_ω*) where *D_ab_ω* is the phase shift due to a transmission delay. To simplify notations, we let Φ_*a*_ = *ϕ_a_* + *ϕ_s_* be the sum of phase shifts due to population response of *a* and synaptic dynamics (without a delay), and Φ_*ab*_ = Φ_*a*_ + Φ_*b*_ etc.

The real part of Eq. 11 is

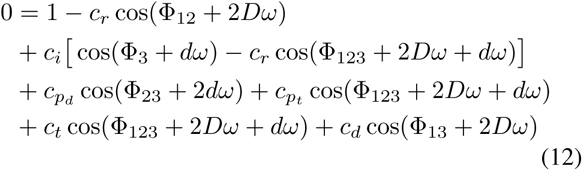

and its imaginary part is

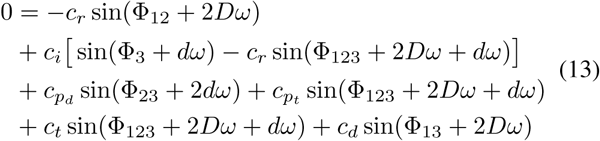

where 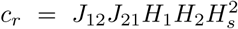 is the bidirectional coupling between *E*_1_ and *E*_2_, *c_i_* = *J*_33_*H*_3_*H_s_* is the *I*_3_-*I*_3_ coupling, 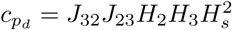 and 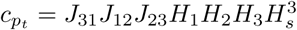 are the disynaptic and trisynaptic *E*_2_-*I*_3_ couplings, respectively, and 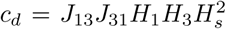 and 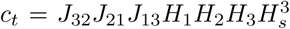 and the disynaptic and trisynaptic *E*_1_-*I*_3_ couplings, respectively.

In the following calculations, *J*_33_ and *D* are the two bifurcation parameters, and we sought to express them as functions of *ω* (See Appendix A for the derivation of other critical couplings). First, to write *J*_33_ as a function of *ω*, we removed *D* from Eqs. 12 and 13 by moving three (two) terms in Eq. 12 (Eq. 13) that did not include *D* to the other side of the equation, squaring both sides of each equation, then adding two equations to obtain a quadratic equation of *c_i_*,

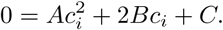

Then,

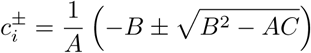

or

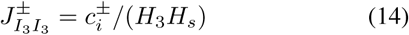

where

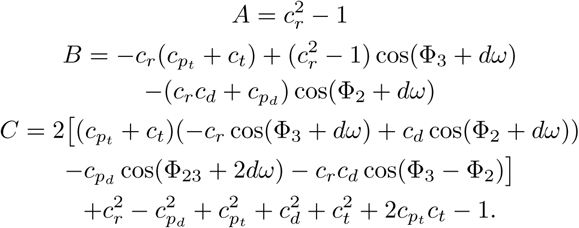

Next, to write *D* as a function of *ω*, we invoked trigonometric identities in Eqs. 12 and 13 to derive a system of equations to explicitly solve for cos 2*Dω* and sin 2*Dω*.

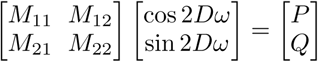

where

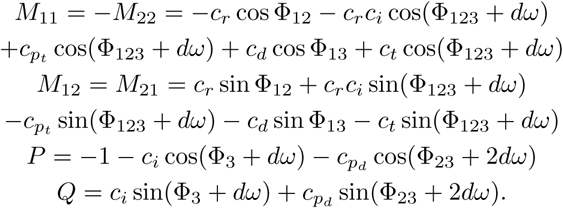

Substituting 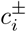 calculated above, we obtained an expression for the lateral delay for Hopf bifurcation

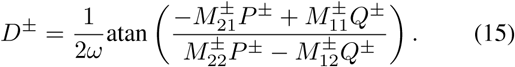

In the following sections, we describe the Hopf bifurcation lines 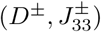 that satisfy both Eqs. 13 and 12 by varying *ω* and compare the analytical results with numerical solutions of delay differential equations for the rate model and simulations of networks of leaky integrate-and-fire neurons for the spiking network model.

## IV. DYNAMICAL STATES OF *E*_1_*E*_2_*I*_3_ NETWORK

In the following, we refer to the (non-oscillatory) steadystate as *S* and three oscillatory states as *O*_1_, *O*_2_ and *O*_3_. As described in Eq. 11, three types of inhibitory couplings *J*_13_, *J*_23_ and *J*_33_ are responsible for generating the oscillatory states *O*_1_, *O*_2_ and *O*_3_, respectively.

### A. Network activity states

To characterize different dynamical states of the three population network, we systematically varied the *I*_3_-*I*_3_ coupling (*J*_33_) and the delay in lateral connections (*D*) (Fig 2). For each parameter pair (*J*_33_, *D*), we simulated the network for 1.2 seconds and measured the standard deviation of the population rates to estimate whether the network exhibited oscillations. The external input to each population was adjusted to maintain constant population rates across different network setups (5 Hz for excitatory and 10 Hz for inhibitory spiking neurons) while other network parameters remained fixed. For both network models, the standard deviation of each population rate was normalized by its means and averaged over three populations to obtain the coefficient of variation of the population activity. When the network was oscillating, the standard deviation of the network activity was higher as the network activity waxed and waned. This is, however, only an indirect measure and may not directly imply oscillations, therefore, we also examined the spectrum of the population activity (Fig. 3).

**FIG. 2.**
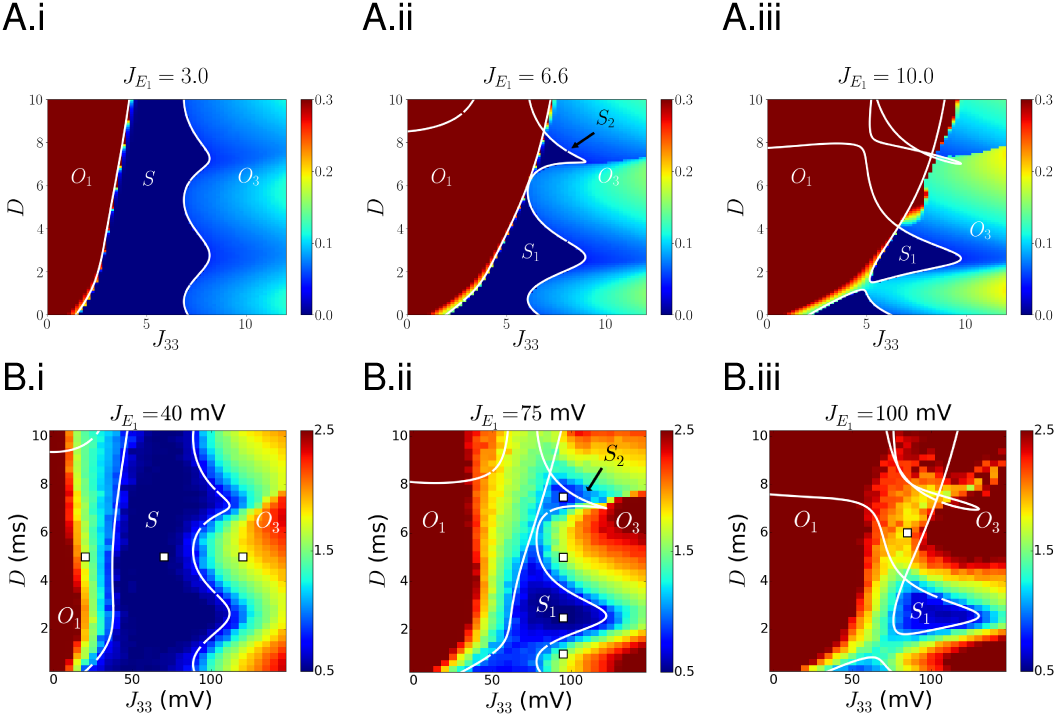
Bifurcation diagrams of (**A**) rate model and (**B**) spiking network model as a function of *I*_3_-*I*_3_ coupling *J*_33_ and lateral delay *D*. The strength of synaptic projection, 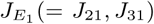, from *E*_1_ to *E*_2_*I*_3_ is weak (A.i, B.i), intermediate (A.ii, B.ii) and strong (A.iii, B.iii). For the rate model, *J*_12_ = 0.5, *J*_13_, *J*_23_, *J*_32_ = 2, *d* = 2.5, *τ_d_* = 0. For the spiking network model, the total (individual) synaptic weights *J*_12_ = 30(0.03), *J*_13_ = 80(0.32), *J*_23_, *J*_32_ = 50(0.2, 0.05) mV. Note that the weights of individual synapses from neuron *i* in population *a* to neuron *j* in population *b* is given by *J_ab_*/(*pN_b_*) (Eq. 3), so that *I*_3_ to *I*_3_ synaptic weights are between 0 and 0.6mV. Local delay *d* = 2.5ms, and synaptic decay time *τ_d_* = 1ms. Color bars show the coefficient of variations of the network models (See text for details). White lines show analytical estimates of a Hopf bifurcation (Eqs. 14 and 15).

**FIG. 3.**
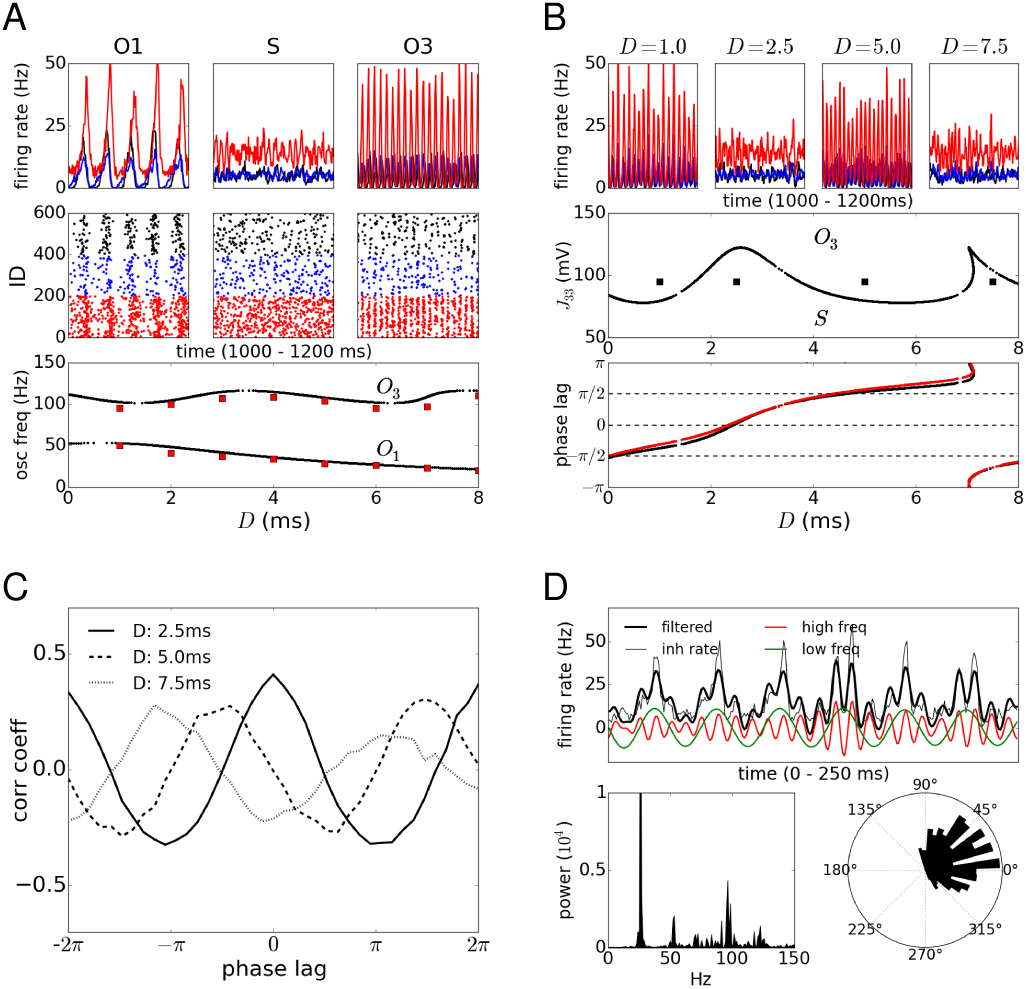
(**A**) (top) Instantaneous firing rate of *E*_1_ (black), *E*_2_ (blue) and *I*_3_ (red) populations in *S, O*_1_ and *O*_3_ states; (middle) Spike raster of the corresponding spiking activity. (bottom) Oscillation frequency of *O*_1_ and *O*_3_ states as a function of lateral delay; black: analysis, red: simulations (**B**) (top) Alternation between *S* and *O*_3_ states due to the lateral delay, corresponding to white squares in Fig. 2B.ii. (middle) Analytical estimates of critical 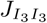 (black line) and network states corresponding to the top row (black squares). (bottom) Analytical estimate of the phase lag Δ*ϕ*_31_ of *I*_3_ with respect to *E*_1_ in the *O*_3_ state; black: full network (Eq. (B1)), red: simplified *E*_1_*I*_3_ network (Eq. (19)). (**C**) Cross-correlation of mean firing rate of *E*_1_ and *I*_3_, calculated using simulation results from part (B). (**D**) Cross-frequency coupling appears when *O*_1_ and *O*_3_ merge; (top) mean firing rate of *I*_3_ (gray), its high (red) and low (green) frequency components, and the sum of high and low frequencies (black); (left bottom) power spectrum of *I*_3_ mean firing rate; (right bottom) Correlation between fast oscillation amplitude (black bar) and slow oscillation phase (angle).

For a fixed value of *D*, the network exhibited three distinct states, *O*_1_, *S* and *O*_3_, as we varied *J*_33_. (Hopf bifurcation estimates are shown as white lines in Fig. 2). The estimates of the network activity states closely matched with the states obtained from a corresponding simulations of a network of spiking neurons (Figs. 2A, B). For low values of *J*_33_, *O*_1_ was observed which is characterized by slow frequency oscillations (≈ 25 Hz) mediated by the *E*_1_-*I*_3_ loop with lateral delay *D* = 5 ms (see white squares in Fig. 2B.i and the corresponding spiking activity in Fig. 3A, top). For moderate values of *J*_33_, oscillations vanished and a non-oscillatory state emerged (*S*) in which neurons fired asynchronously and the population firing rate randomly fluctuated around the steady-state (akin to the asynchronous irregular state of [3]). Finally for very strong inhibitory coupling (high *J*_33_), the non-oscillatory state was transformed into another oscillatory state *O*_3_, which was characterized by high frequency oscillations (≈ 100 Hz) mediated by the local *I*_3_-*I*_3_ loop with local delay *d* = 2.5 ms.

As the lateral delays *D* was increased the regions of *O*_1_ monotonically creased and the state *S* was observed at higher values of *J*_33_. The lateral delays *D* had a more dramatic effect on the emergence of the state *O*_3_. The value of *J*_33_ at which the oscillatory state *O*_3_ was observed varied in a periodic manner as *D* was increased (see the interface of *O*_1_ and *S* and that of *O*_3_ and *S* in Figs. 2A.i and B.i). While, the oscillation frequency in *O*_1_ state decreased monotonically with *D*, the oscillation frequency in *O*_3_ state was independent of *D* (Fig 3A, bottom; black lines: analysis, red squares: simulations).

To better understand why the critical *J*_33_ weights varied in a periodic manner as a function of lateral delays *D*, we next examined the oscillation dynamics at the interface of the nonoscillatory state *S* and the oscillatory state *O*_3_

### B. Modulation of critical *I*_3_-*I*_3_ coupling due to lateral delay

To gain intuition on why the local coupling *J*_33_ may depend on lateral delays *D*, we reduced the subnetwork *E*_2_*I*_3_ to an inhibitory population *I*_3_. This effectively assumes that the subnetwork *E*_2_*I*_3_ operates in an inhibition dominated regime. By solving Eqs. 12 and 13 for the critical *I*_3_-*I*_3_ coupling in terms of *D* we obtained

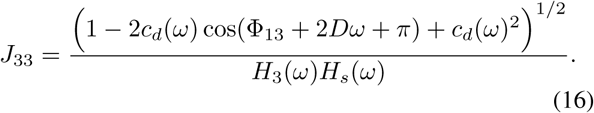

From Eq. 16, the reduced network model suggests that the critical *J*_33_ modulates periodically as a function of *D*, and the period of modulation *T* = *π/ω*\* = 1/(2*f*\*) is determined by the frequency of network oscillations, *ω*\* = 2*π*\*. (Here we assumed that the oscillation frequency at the transition to *O*_3_- state depends weakly on *D*; in other words, *ω*(*D*) ≡ *ω*\* on the boundary of *O*_3_ as shown in Fig. 3A (bottom).) Moreover, the maximum of *J*_33_ is attained at

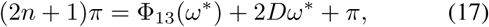

and the minimum of *J*_33_ at

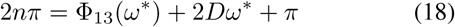

for *n* = 0, ±1, …, where Φ_13_(*ω*\*) + 2*Dω*\* is the total phase shift induced by the bidirectional lateral connections between *E*_1_ and *I*_3_, and *π* appears due to the inhibitory coupling. In other words, Eq. 17 (Eq. 18) suggests that the critical *J*_33_ reaches its maximum (minimum) when the network oscillation relayed through the lateral connections is anti-phase (in-phase) to the oscillations generated in *I*_3_.

Conceptually, this phenomenon can be understood as follows. If oscillatory activity relayed through lateral connections is in-phase with oscillatory activity generated by the local *E*_2_-*I*_3_ loop, weak coupling is sufficient to induce network-wide oscillations. On the other hand, if the relayed network activity is anti-phasic to locally generated activity, stronger coupling is required to overcome the suppression of the local activity by the relayed activity. Because changing the lateral delay shifts the phase of relayed activity continuously, the critical coupling strength modulates qusi-periodically with a period determined by the oscillation frequency. In Fig. 2B.i, for instance, the period of *O*_3_-boundary (*T* ≈ 5 ms) is determined by the oscillation frequency of *O*_3_ state (*f* ≈ 100 Hz, Fig. 3A, bottom): *T* ≈ 1/(2*f*) as predicted by Eq. 16.

We also calculated the relative phase 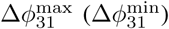 of *I*_3_ population rate with respect to the *E*_1_ population rate at the maximum (minimum) of *J*_33_. For the reduced network model, the phase lag is given by

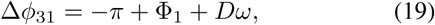

and at the max and min of *J*_33_,

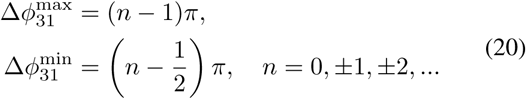

where we used Eqs. 17 and 18 to evaluate *Dω*\* and assumed that the phase shifts due to population response functions are equal, i.e. Φ_1_ = Φ_3_. We then verified numerically that the phase lag in the full *E*_1_*E*_2_*I*_3_ network can be approximated by that of the reduced network (Fig. 3B, bottom; black: full network, red: reduced network). See Appendix B for the derivation of phase lags in the full and reduced network models.

### C. Cross-frequency coupling

The network architecture investigated here has been suggested to generate cross-frequency oscillations in which power in a frequency band is modulated by the phase of another oscillation [13]. In our model, multi-frequency oscillations appeared as we increased the strength of long range excitation 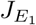.

At an intermediate step towards the emergence of multifrequency oscillations, we examined how the strength of connections from the excitatory population 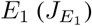 affected the landscape of the network states. First, we found that increase in 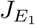 increased the region of the state *O*_1_. Second, 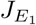 also increased the modulation in the boundary of state *O*_3_. Together, these changes meant that the region of non-oscillatory state was not observed for some values of *D*, such that the non-oscillatory state appeared in isolated regions, *S*_1_ and *S*_2_ in Figs. 2A.ii and B.ii. There was a wide range of inhibitory coupling *J*_33_ (e.g. 80 - 120 mV in Fig. 2B.ii), over which the network can switch between non-oscillatory and oscillatory states by varying the lateral delay (white squares in Fig. 2B.ii and the corresponding spike activity shown in Fig. 3B, top and middle).

As the strength of synaptic input from *E*_1_ was further increased, two Hopf bifurcation lines defining the *O*_1_ and *O*_3_states merged and created a small region in the space spanned by *J*_33_ and *D* in which the network exhibits non-oscillatory activity *S*_1_ (Fig 2A.iii and B.iii). Outside of *S*_1_, where *O*_1_ and *O*_3_ merged (e.g. white square in Fig. 2B.iii), the slow oscillatory activity induced by the lateral *E*_1_-*I*_3_ loops and the fast oscillatory activity induced by the local *I*_3_-*I*_3_ loop coexisted. This was evident in the power spectrum of inhibitory population firing rate, which showed two peaks at low (25 Hz) and high (100 Hz) frequencies (Fig. 3D, left bottom). Moreover, when the population rates were band-passed filtered at high and slow frequencies then summed up, the filtered population rates closely followed the actual inhibitory firing rates (Fig. 3D, top). Interestingly, in our model the amplitude of fast oscillation was modulated according to the phase of slow oscillation (Fig 3D, right bottom). This is akin to the modulation of gamma band osculations power by the phase of theta oscillations [23] in the hippocampus.

Thus, we show that when a partially overlapping neuron population participates in generation of both fast oscillation and slow oscillation (i.e. *I*_3_ is part of the fast *I*_3_-*I*_3_ loop and the slow *E*_1_-*I*_3_ loop), cross-frequency coupling emerges in which slow oscillation generated via the lateral loop (*E*_1_-*I*_3_) with long delay *D* modulates the amplitude of fast oscillation (*I*_3_-*I*_3_) by providing periodic input.

### D. Emergence of an oscillatory state driven by *E*_2_-*I*_3_ coupling

Thus far, we considered oscillatory activity generated by the *I*_3_-*I*_3_ loop. In this section, we describe how including the oscillatory activity due to the *E*_2_-*I*_3_ loop further enriches the dynamical landscape of the three population network. To control the interaction of the *E*_2_-*I*_3_ and *I*_3_-*I*_3_ loops, we introduce a parameter that determines the relative strength of the *E*_2_-*I*_3_ coupling with respect to *I*_3_-*I*_3_ coupling: *J*_23_ = *αJ_I_, J*_33_ = *J_I_*.

With weak coupling between *E*_2_-*I*_3_ (i.e. small *α*), Hopf bifurcation structure was identical to the one obtained by varying *J*_33_ alone, as shown in Figs. 2A.i and 2B.i. In other words, the *O*_3_ state, driven by the *I*_3_-*I*_3_ loop, was the only oscillatory state that can be generated by the *E*_2_*I*_3_ subnetwork. When *α* was increased, a new oscillatory state *O*_2_, driven by the *E*_2_-*I*_3_ loop, emerged in the small isolated regions in the space spanned by *J*_1_ and *D* (Figs. 4A.i and B.i). The *O*_2_ state emerged in a parameter space which led to the non-oscillatory state (*S*) for low values of *α*. That is, for high *α* values, long-range interactions with the population *E*_1_ destabilized the non-oscillatory state of the *E*_2_*I*_3_ subnetwork to create oscillations.

**FIG. 4.**
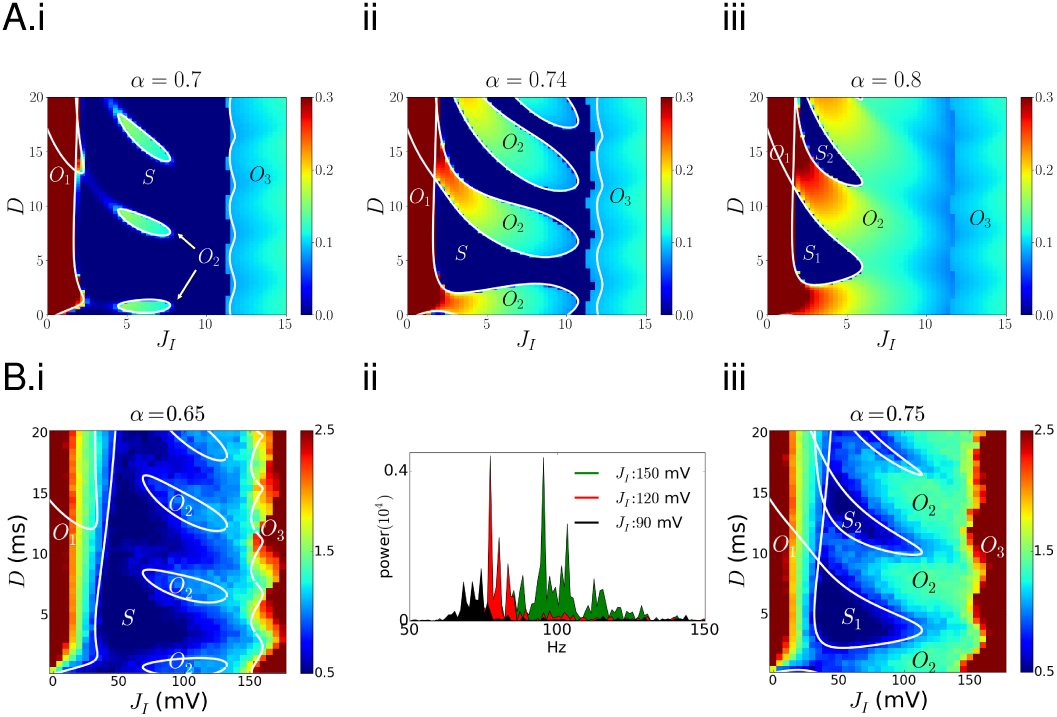
Composition of three oscillatory states. denotes the relative strength of *E*_2_-*I*_3_ loop: *J*_23_ = *J_I_, J*_33_ = *J_I_* (**A**) Rate model; *J*_12_ = 0.5, *J*_13_ = 2, *J*_32_ = 4, *J*_21_ = *J*_31_ = 1, *d* = 2.5, *τ_d_* = 0. (**B**) Spiking network model; B.ii, Power spectrum of population *I*_3_’s mean firing rates for *D* = 7ms in B.i.; Total (individual) synaptic weights, *J*_12_ = 30(0.03), *J*_13_ = 60(0.24), *J*_32_ = 120(0.12), *J*_21_ = *J*_31_ = 30(0.06) mV, *d* = 2.5 ms, *τ_d_* = 1 ms. Color bars show the coefficient of variations of network models. White lines show analytical estimates of a Hopf bifurcation (Eqs. A2 and 15).

To better understand why non-oscillatory and different oscillatory states appear as a function of *J_I_*, we fixed the lateral delay (e.g. *D* = 9 in Fig. 4A.i) and examined the changes in network state as *J_I_* increases. For small *J_I_*, local inhibition was too weak to withstand the oscillatory instability driven by the lateral *E*_1_-*I*_3_ loop, so the network entered the *O*_1_-state. The network shifted to the non-oscillatory state *S* when the local inhibition *J_I_* was increased. When *J_I_* became sufficiently strong to generate oscillatory activity through the *E*_2_-*I*_3_ loop, the network entered *O*_2_-state. When *J_I_* was further increased, the increased *I*_3_-*I*_3_ coupling suppressed the oscillatory activity induced by the *E*_2_-*I*_3_ loop, which brought the network back to the non-oscillatory state. For strong *J_I_*, the *I*_3_-*I*_3_ coupling becomes the dominant network motif producing oscillatory activity, hence the network entered the *O*_3_-state.

The *O*_2_-state, however, appeared only within a restricted range of lateral delays (e.g. 7 < *D* <10 in Fig. 4A.i) and vanished gradually outside of this range. As discussed in Section IV B, the *O*_2_ state appears when the oscillatory activity relayed through the lateral connections is in-phase with the ongoing oscillations in the *E*_2_*I*_3_ subnetwork. On the other hand, the *O*_2_ state can no longer exist when the relayed network activity no longer enhances the local activity. Such resonance and cancellation effects, occurring repeatedly, gave rise to isolated *O*_2_-states at multiple sites. The periodic appearance of stationary and oscillatory states is a robust phenomenon in delayed feedback system, which has been investigated in the context of controlling pathological brain rhythms [18, 20].

When the *E*_2_-*I*_3_ coupling became stronger (*α* was increased), the *O*_2_ state expanded across the non-oscillatory state *S* and merged with the *O*_1_ state, creating a complex configuration of multiple network states as shown in Fig. 4A.ii. The *O*_1_ and *O*_3_ states, previously separated by *S*, were now bridged by an elongated *O*_2_ region. When *α* was further increased, all three oscillatory states, *O*_1_, *O*_2_ and *O*_3_, appeared contiguously, and the non-oscillatory states formed isolated regions surrounded by the *O*_1_ and *O*_2_ states (Figs. 4A.iii and B.iii). Such dynamical landscape is similar to the previously discussed bifurcation structure that produced cross-frequency oscillations at a parameter region where multiple Hopf bifurcations meet (see Figs. 2 A.iii and B.iii).

In networks of spiking neurons, *O*_2_-states emerged from the non-oscillatory state at multiple sites, as predicted by the analysis. However, unlike the dynamical landscape of the firing rate model (Fig. 4A.i), network simulations showed that *O*_2_ states did not exist in isolation. The network activity remained oscillatory in-between the *O*_2_ and *O*_3_ states (Fig. 4B.i), and the oscillation frequency increased gradually as the network transitioned from *O*_2_, *S*, to *O*_3_ (Fig 4B.ii). When became large as shown in Fig. 4B.iii, the *O*_2_ state expanded horizontally across the non-oscillatory state, connected the *O*_1_ and *O*_3_ states, and created isolated non-oscillatory states surrounded by the oscillatory states, *O*_1_ and *O*_2_, similarly to the firing rate model (Fig. 4A.iii).

## V. CONCLUSION

Population of neurons embedded in a larger network rarely acts alone but interacts in concert with neighboring neurons to produce network activity. In this study we extended the standard two-population model consisting of excitatory and inhibitory neurons, and demonstrated in a minimal three-population model that when an excitatory-inhibitory network is coupled to an additional excitatory population, the lateral connections between them can create a rich bifurcation structure, composed of isolated stationary states, multiple oscillatory states and cross-frequency coupling. Such dynamical landscape allows the network to fix its operating point in the non-oscillatory stationary state and easily tap into various oscillatory states generating slow frequency oscillations (via lateral *E*_1_-*I*_3_ loop), fast frequency oscillations (via local *E*_2_-*I*_3_ and *I*_3_-*I*_3_ loops), or nested (cross-frequency) oscillations (by positioning itself in a region where slow and fast oscillations merge). We also found that two types of local oscillations (*O*_2_ and *O*_3_) can be interwoven intricately with the stationary states, which has not been observed in isolated excitatory-inhibitory networks.

Previously, the coupling of gamma (30-80 Hz) and theta (4-12 Hz) frequency oscillations in the hippocampus was investigated using a network model composed of one excitatory and two types of inhibitory neuron models, where one of the inhibitory neuron type (O-LM interneuron) was responsible for generating the gamma rhythm [24]. Our results, as suggested in [13], demonstrate a different mechanistic model for cross-frequency coupling where there is only one type of inhibitory neuron but two modes of oscillations can be generated via long lateral and short local delays.

Various effects of time delay on neural dynamics have been studied extensively in neural field equation that models spatial interactions. It has been shown that the delay can induce oscillations for local excitation-lateral inhibition interaction [25], give rise to rich bifurcation structure in simple scalar model [26], and stabilize stochastic bump attractors [27].

Our results demonstrate that a lumped firing rate model and randomly connected spiking models can also develop rich dynamic repertoire without any spatial interactions.

To cleary expose the rich dynamics of the three population network we have used rather wide range of delays (0–20 ms). Long delays beyond 10 ms are usually not observed in biological neuronal networks. All the key dynamical states (*O*_1_, *S, S*_1_, *O*_2_, *O*_3_ can be observed for lateral delays up to 5 ms which are known to exist in the brain. For instance, the thalamocortical loop has 3.4 ms delay in mice [28]) and the inter-hemispheric connections can show delays of 3 – 9 ms in different species [29]. In primates inter-hemispheric delays are about 4 ms [30]. Our results show that slow conductation delays in thalamocortical loops and inter-hemispheric connections can play a very important role in reshaping the local network dynamics and introducing new dynamics through the long range interactions. Our network simulations and analysis was restricted to a special case in which all delays were fixed. It would be of interest to study the effects of distributed delays in the future, as in [31–33], to reflect biologically realistic connectivity and go beyond the three population motifs.

## ACKNOWLEDGMENTS

This work was supported by the BrainLinks-BrainTools Cluster of Excellence funded by the German Research Foundation (DFG EXC 1086), the German Federal Ministry of Education and Research (FKZ 01GQ0830), and the INTERREG IV Rhin superieur program and European Funds for Regional Development through the project TIGER A31.

## Appendix A: Calculation of critical couplings

We substitute *λ* = *iω* and calculate the rate response function *R_a_*(*iω*) and the synaptic kernel ∑_*ab*_(*iω*). For the rate model,

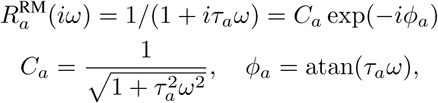

with 0 ≤ atan(*τ_a_ω*) < *π*/2, and for the spiking network,

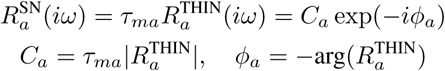

where 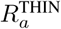 is the population response function obtained numerically from the threshold integration method [22]. The synaptic kernel

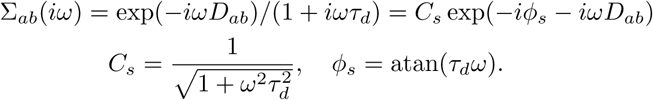

with 0 ≤ atan(*τ_d_ ω*) < *π*/2.

Substituting *R_a_*(*iω*) and ∑_*ab*_(*i ω*) to Eq. (11) yields

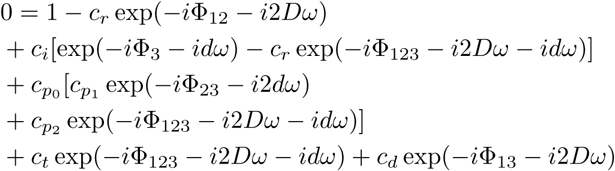

where we decompose 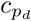 and 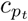 in Eqs. (12) and (13) in terms of 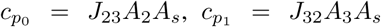 and 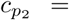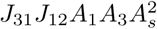 in order to explicitly solve for *J*_23_ in the following calculations.

To study the bifurcaiton structure when *J*_23_ are varied, we manipulated Eqs. (12) and (13), as discussed in Section III B, to solve for the inhibitory coupling strength *J*_23_ = 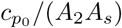.

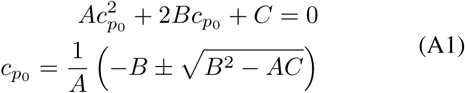

where

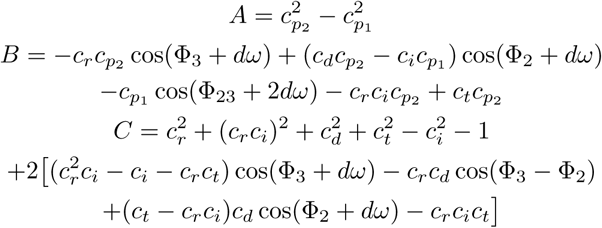

On the other hand, to study the bifurcation structure when both *J*_23_ and *J*_33_ are varied, we similarly manipulate Eqs. (12) and (13) to solve for the inhibitory coupling *J_I_* where *J*_23_ = *J_I_* and *J*_33_ = *J_I_*.

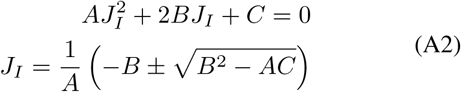

where

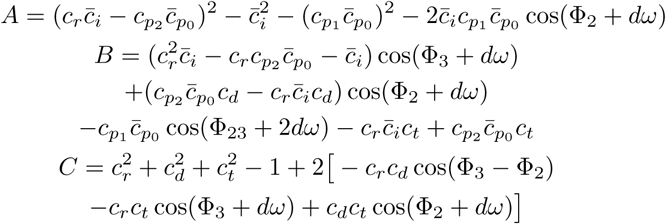

and 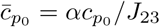 and 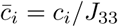.

## Appendix B: Phase lag

From the second and third lines of the coefficient matrix, Eq. (10), we can derive

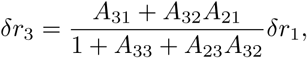

which implies that the relative phase difference

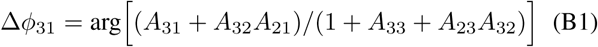

of *I*_3_ with respect to *E*_1_ is determined by the later couplings, *A*_31_ and *A*_32_*A*_21_, that connect *E*_1_ to *E*_3_, and the local couplings, *A*_33_ and *A*_23_*A*_32_. In Fig 3B, we numerically evaluate the above Δ*ϕ*_31_.

On the other hand, for the reduced two-population network considered in Section IV B, the first line of Eq. (10),

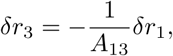

gives a simple expression for the phase difference

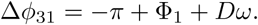

